# Single position substitution of hairpin pyrrole-imidazole polyamides imparts distinct DNA-binding profiles across the human genome

**DOI:** 10.1101/2020.08.13.249730

**Authors:** Paul B. Finn, Devesh Bhimsaria, Asfa Ali, Asuka Eguchi, Aseem Z. Ansari, Peter B. Dervan

**Author notes:** Department of Bioengineering, Stanford University, Stanford, California, United States of America (Paul B. Finn). Corresponding author (PBD). These authors contributed equally to this work (Paul B. Finn, Devesh Bhimsaria).

## Abstract

Regulating desired loci in the genome with sequence-specific DNA-binding molecules is a major goal for the development of precision medicine. Pyrrole–imidazole (Py–Im) polyamides are synthetic molecules that can be rationally designed to target specific DNA sequences to both disrupt and recruit transcriptional machinery. While *in vitro* binding has been extensively studied, *in vivo* effects are often difficult to predict using current models of DNA binding. Determining the impact of genomic architecture and the local chromatin landscape on polyamide-DNA sequence specificity remains an unresolved question that impedes their effective deployment *in vivo*. In this report we identified polyamide–DNA interaction sites across the entire genome, by covalently crosslinking and capturing these events in the nuclei of human LNCaP cells. This method, termed COSMIC-seq, confirms the ability of hairpin-polyamides, with similar architectures but differing at a single ring position, to retain *in vitro* specificities and display distinct genome-wide binding profiles. These results underpin the development of Py-Im polyamides as DNA-targeting molecules that mediate their regulatory or remedial functions at desired genomic loci.

## INTRODUCTION

Regulating genomic architecture and activity with sequence-specific synthetic DNA binding molecules is a long-standing goal at the interface of chemistry, biology and medicine. Small molecules that selectively target desired genomic loci could be harnessed to regulate critical gene networks. The greatest success in designing small molecules with programmable DNA-binding specificity has been with pyrrole-imidazole (Py-Im) polyamides [1–8]. Pyrrole-imidazole (Py-Im) polyamides are synthetic DNA-binding oligomers with high sequence specificity and affinity [7]. An oligomer, comprising a modular set of aromatic pyrrole and imidazole amino acids linked in series by a central aliphatic γ-aminobutyric acid (GABA) ‘turn’ unit, fold into a hairpin structure in the minor groove of DNA and afford binding affinities and specificities comparable to natural transcription factors [3,7]. Sequence specificity is programmed through side-by-side pairs of the Py and Im subunits that “read” the steric and hydrogen bonding patterns presented by the edges of the four Watson-Crick base pairs on the floor of the minor groove [5]. DNase I footprinting titrations and other *in vitro* methods have extensively characterized the binding affinity and specificity of these molecules [3,6,7,9]. An Im/Py pair binds G•C; Py/Im binds C•G, and Py/Py pairs both bind A•T and T•A (denoted as W) [1,2]. Py-Im polyamide binding in the minor groove induces allosteric changes to DNA, widening the minor groove and narrowing the major groove [10–12]. Polyamide-DNA binding is sufficient to disrupt protein-DNA interfaces, including DNA interactions made by transcription factors and the transcriptional machinery [13–15]. Additionally, polyamides can function as sequence-specific synthetic cofactors through allosteric DNA modulation to enhance the assembly of protein-DNA complexes [12]. Py-Im polyamides are cell permeable, localize to the nucleus in live cells and are non-genotoxic [16–18] failing to activate canonical DNA damage response or significantly alter cell cycle distribution [19].

The identification of new mechanistic insights into Py-Im polyamide activity have underlined the importance of mapping polyamide binding to chromatin [15,18,19]. Polyamide binding in the more complex cellular environment presents a formidable challenge since chromatin DNA has varying degrees of accessibility. Sequence specific access by Py-Im polyamides to the nucleosome core particle (NCP) has been demonstrated *in vitro* and with x-ray crystal structures of NCP•polyamide complexes [20–22]. However, the extent to which chromatin states influence polyamide binding to its cognate sites remains a long-standing question. The lack of clarity on the parameters that govern genome-wide binding of polyamides greatly impedes the deployment of this class of molecules to regulate cell fate-defining and disease-causing gene networks *in vivo*.

We report here the genome-wide binding profiles of two Py-Im polyamides **1** and **2**, of identical architecture (8-ring hairpin) that differ at a single aromatic ring position in cellular nuclei using COSMIC-seq (‘crosslinking of small molecules for isolation of chromatin with next-generation sequencing), **Fig 1** [23,24]. COSMIC-seq employs a tripartite conjugate composed of the DNA-binding ligand attached to a biotin affinity handle and a psoralen photocrosslinker. Genome-wide binding of these tripartite molecules is captured by photo-induced crosslinking followed by biotin-enabled enrichment and unbiased NGS sequencing of the conjugated genomic loci [23,24]. The ability to induce rapid crosslinking at the desired time point distinguishes COSMIC-seq from continuous and uncontrolled alkylation-dependent DNA conjugations that have been used to query genome-wide binding of polyamides [25]. COSMIC-seq also differs from Chem-seq approaches that use ligands for protein complexes that are associated with the genome [26]. Previously, COSMIC-seq was utilized to access genome-wide binding of two structurally distinct Py-Im polyamides (hairpin vs linear) that code for very different sequences [24]. An 8-ring hairpin Py-Im polyamide (TpPyPyIm-γ-PyImPyPy-β-Dp) binds 6 bp of DNA (5’-WTWCGW-3’) [27], whereas a linear polyamide (ImPy-β-ImPy-β-Im-β-Dp) binds 9 bp of purine rich DNA (5’-AAGAAGAAG-3’) [28–31]. While such a dramatic difference in target sequence composition leads to distinct genome-wide binding profiles, we wondered how a more challenging single position change (CH to N:) within one ring of an 8-ring hairpin would affect genomic occupancy. In this study we applied COSMIC-seq to determine if two polyamides of identical size and architecture, hairpins **1** and **2** which code for 6 base pair sites differing by one base pair position 5’-WGWWCW-3’ and 5’-WGGWCW-3’, respectively, can display distinct genomic binding occupancy on chromatin. These experiments provide a more stringent test of genome-wide binding properties of hairpin polyamides in a chromatin environment for application as precision-targeting molecules.

**Fig 1.**
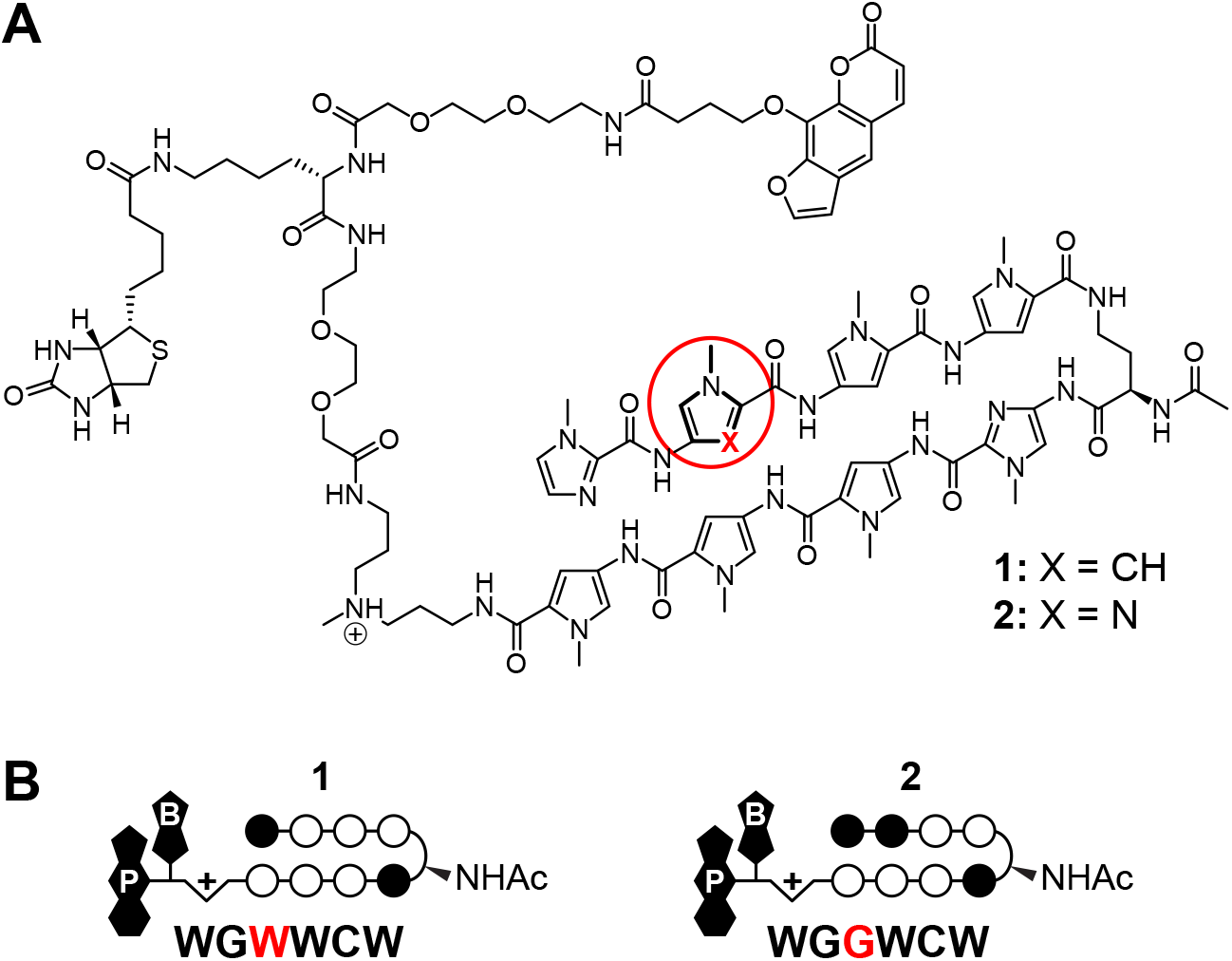
Trifunctional Py-Im polyamide conjugates 1 and 2. (**A**) Chemical structure of hairpin Py-Im polyamides **1** and **2** which differ by one atom, shown in red, and (**B**) the corresponding predicted target sequences based on the pairing rules. Py-Im polyamide **1** targets the DNA sequence 5’-WGWWCW-3’ and Py-Im polyamide **2** targets 5’-WGGWCW-3’. Open and filled circles represent N-methylpyrrole (Py) and N-methylimidazole (Im), respectively. The N-acetylated (R)-γ-aminobutyric acid turn residue is shown as a semicircle, and psoralen and biotin are denoted by P and B, respectively.

## MATERIALS AND METHODS

### Materials

Chemicals and solvents were purchased from standard chemical suppliers and used without further purification. (R)-2,4-Fmoc-Dab(Boc)-OH (α-amino-GABA turn) was purchased from Peptides International. Monomers were synthesized as previously described [32]. Kaiser oxime resin (100-200 mesh) and benzotriazole-1-yl-oxy-trispyrrolidinophosphonium hexafluorophosphate (PyBOP) were purchased from Novabiochem. 2-Chlorotrityl chloride resin was purchased from Aapptec. Preparative HPLC purification was performed on an Agilent 1200 Series instrument equipped with a Phenomenex Gemini preparative column (250 × 21.2 mm, 5μm) with the mobile phase consisting of a gradient of acetonitrile (CH_3_CN) in 0.1% aqueous trifluoroacetic acid (TFA). Polyamide concentrations were measured by UV/Vis spectroscopy in distilled and deionized water (ddH_2_O) with a molar extinction coefficient of 8650 M^−1^ cm^−1^ at 310 nm for each *N*-methylpyrrole (Py) and *N*-methylimidazole (Im) and 11,800 M^−1^ cm^−1^ for the psoralen/biotin derivative **3** [33,34]. Analytical HPLC analysis was conducted on a Beckman Gold instrument equipped with a Phenomenex Gemini analytical column (250 × 4.6 mm, 5μm), a diode array detector, and the mobile phase consisting of a gradient of acetonitrile in 0.1% aqueous TFA. Matrix-assisted, LASER desorption/ionization time-of-flight (MALDI-TOF) mass spectrometry was performed on an Autoflex MALDI TOF/TOF (Bruker) using α-cyano-4-hydroxycinnamic acid matrix. Oligonucleotides were purchased from Integrated DNA Technologies Inc. All sequencing samples were processed as single read (50 bp) sequencing runs at the California Institute of Technology Millard and Muriel Jacobs Genetics and Genomics Laboratory on an Illumina HiSeq 2500 Genome Analyzer.

### Chemical synthesis

Polyamides **1A** and **2A** were synthesized on solid support (Kaiser oxime resin, 100-200 mesh), using microwave-assisted PyBOP coupling conditions with *N*-methylpyrrole (Py), *N*-methylimidazole (Im) amino acid monomers and dimers (**5a** & **5b**) as previously described, **S1A Fig** [34]. Polyamides were cleaved from resin with neat 3,3′-diamino-*N*-methyldipropylamine (60 °C, 5 min, μW), precipitated with diethyl ether at −20 °C, re-dissolved in 20 - 30% (v/v) CH_3_CN/H_2_O (0.1% TFA), and purified by reverse-phase preparative HPLC. Fractions that showed clean polyamide without contaminants were frozen in liquid nitrogen and lyophilized to dryness as a white-yellow solid. The identity and purity were confirmed by MALDI-TOF mass spectrometry and analytical HPLC. The observed mass for **1A** (C_59_H_75_N_22_O_10_) is 1251.78 (calculated 1251.60) and for **2A** (C_58_H_74_N_23_O_10_) is 1252.75 (calculated 1252.60).

The psoralen-biotin peptide **3** was synthesized by manual Fmoc solid-phase synthesis on 2-chlorotrityl chloride resin by standard procedures, **S1B Fig** [23]. Coupling and deprotection were performed at room temperature for 1 h and 15 min, respectively. Briefly, Fmoc-protected amino acids or polyethylene glycol (PEG) linkers were activated with HATU and HOAt in the presence of *N*, *N*-diisopropylethylamine (DIPEA) in dimethylformamide (DMF) (or DMSO/DMF) and deprotection of the Fmoc group was achieved with 20% piperidine in DMF. Cleavage from resin was achieved with a solution of 95% (v/v) TFA, 2.5% (v/v) H_2_O, and 2.5% (v/v) triisopropylsilane and purified by reverse-phase preparative HPLC, lyophilized to dryness as a white powder and protected from light. The identity and purity were confirmed by MALDI-TOF mass spectrometry and analytical HPLC. The observed mass for **3** (C_43_H_61_N_6_O_15_S) is 933.36 (calculated 933.39).

Polyamide-peptide conjugates **1** and **2** were synthesized by solution phase peptide coupling conditions and protected from light, **S1C Fig**. Peptide acid **3** (1 equiv.) was pre-activated for 5 min at room temperature with a solution of HATU/HOAt/DIPEA (3:3:6 equiv.) in DMF. Polyamide **1A** or **2A** was added (1 - 1.5 equiv.), and the coupling was allowed to proceed for 30 - 60 minutes until all of **3** was consumed as determined by analytical HPLC. The polyamide-peptide conjugates were purified by reverse-phase HPLC and lyophilized to dryness. The identity and purity were confirmed by MALDI-TOF mass spectrometry and analytical HPLC. The observed mass for **1** (C_102_H_133_N_28_O_24_S) is 2166.26 (calculated 2165.98) and for **2** (C_101_H_132_N_29_O_24_S) is 2167.23 (calculated 2166.97).

### Cognate site identification

High-throughput cognate binding sites were identified for the polyamides **1** and **2** using SELEX method [35]. A DNA library with a central randomized 20-bp region and flanked by constant sequences (~10^12^ possible sequences, Integrated DNA Technologies) was used for PCR amplification. Polyamide conjugates **1** and **2** at a range of concentrations (5 nM and 50 nM) were added to 100 nM of DNA library in binding buffer [1× PBS (pH 7.6), 50 ng/ μL poly(dI-dC)] and incubated for 1 h at room temperature. Enrichment of the compound-DNA complexes was performed using streptavidin-coated magnetic beads (Dynabeads, Invitrogen) following manufacturer’s protocol. To remove unbound DNA, three washes were done after the capture, with 100 μL ice-cold binding buffer. Beads were resuspended in PCR master mix (EconoTaq PLUS 2× Master Mix, Lucigen), the DNA was amplified for 15 cycles and purified (QIAGEN). Three rounds of selection were performed (DNA was quantified by absorbance at 260 nm before each round of binding). An additional round of PCR was performed after completion of three rounds of selection, to incorporate Illumina sequencing adapters and a unique 6-bp barcode for multiplexing. The starting library was also barcoded and sequenced. Samples were sequenced on an Illumina HiSeq 2500 at the Millard and Muriel Jacobs Genetics and Genomics Laboratory in California Institute of Technology.

### Cell culture conditions

LNCaP cells were maintained in RPMI 1640 (Invitrogen) with 10% Fetal Bovine Serum (FBS, Irvine Scientific) at 37°C under 5% CO_2_. LNCaP cells were purchased from ATCC (Manassas, VA, USA).

### Crosslinking of small molecules for isolation of chromatin with next-generation sequencing

COSMIC-seq was performed in LNCaP nuclei, as previously described [23]. LNCaP cells (~2.5 ×10^7^) were washed twice with cold PBS then resuspended in cold lysis buffer (RSB + 0.1% IGEPAL CA-630, 2.5 ×10^7^ cells/250 μL), incubated on ice for 5 min then centrifuged immediately at 130 × g for 10 min at 4 °C. Nuclei were resuspended in binding buffer [10 mM Tris HCl (pH 8.0), 5 mM MgCl_2_, 1 mM DTT, 0.3 M KCl, 0.1 M PMSF, 0.1 M benzamidine, 0.1 M pepstatin A, 10% glycerol] and treated with psoralen-biotin conjugated polyamide **1** or **2** (0.4 μM and 4 μM, 0.1% DMSO final concentration) for 1 h at 4 °C in the dark. Nuclei were irradiated for 30 min with a UV lamp (2.4 μW/cm^2^; CalSun) through a Pyrex filter, centrifuged at 500 × g and re-suspended in COSMIC buffer [20 mM Tris·Cl (pH 8.1), 2 mM EDTA, 150 mM NaCl, 1mM PMSF, 1mM benzamidine, 1.5 μM pepstatin,1% Triton X-100, 0.1% SDS]. Samples were sonicated at 3 °C for 36 min with a cycle of 10 s ON and 10 s OFF, at HIGH setting (Bioruptor Plus, Diagenode). Samples were centrifuged 10 min at 12,000 × g and 10% of the sample was saved as input DNA and stored at −80 °C until reversal of cross-linking. The rest of the sample was used for the affinity purification (AP). Streptavidin-coated magnetic beads (100 μL per sample, Dynabeads MyOne C1) were washed in COSMIC buffer and incubated with AP samples for 16 h at 4°C. All washes were performed at room temperature unless otherwise noted. For **1** and **2**, **1A** and **2A** were added (5 μM), respectively, in the washes. Samples were washed twice with COSMIC buffer (once 12 h and once 4 h). Samples were then washed once with washing buffer 1 [10 mM Tris·Cl (pH 8.0), 1 mM EDTA, 3% (v/v) SDS], once with washing buffer 2 [10 mM Tris·Cl (pH 8.0), 250 mM LiCl, 1 mM EDTA, 0.5% Nonidet P-40, 1% sodium deoxycholate], twice with freshly prepared washing buffer 3 [4 M urea, 10 mM Tris·Cl (pH 7.5), 1 mM EDTA, 0.1% Nonidet P-40], and twice with TE buffer [10 mM Tris·Cl (pH 8.0), 1 mM EDTA]. Samples were re-suspended in TE and labelled as AP DNA. Input and AP samples were re-suspended in cross-link reversal buffer [10 mM Tris (pH 7.6), 0.4 mM EDTA, 100 mM KOH]. Crosslinks were reversed, and DNA was eluted from beads at the same time by heating samples for 30 min at 90 °C. Input and AP samples were neutralized with 6N HCl, and incubated first with RNase A (0.2 μg/μL) for 1 h at 37 °C and then with Proteinase K (0.2 μg/ μL) for 1 h at 55 °C. Samples were purified with the MinElute PCR Purification Kit (Qiagen).

### CSI data analysis

The reads from Illumina sequencing were de-multiplexed using the 6 bp barcodes and then truncated to include only the 20 bp random portion of the library. On average, 1,031,000 reads per barcode were obtained. The occurrence of every k-mer (8 mer) was counted using a sliding window of size k. To correct for experimental biases and biases in the initial DNA library, a standardized enrichment score was calculated by normalizing the counts of every k-mer from the enriched CSI data (rounds 1, 2 or 3) to the expected number of counts in the library with a fifth-order Markov model derived from the processed library (processed same number of SELEX enrichment rounds without the polyamide as done for the polyamide) [36,37]. The most enriched 8 bp sub-sequences were used to derive position weight matrix (PWM) motifs using MEME [38,39]. Data files for mapped 20 bp reads and normalized 8 bp sequences are available online (https://ansarilab.biochem.wisc.edu/computation.html).

### Sequence logos

PWMs were derived from the 50 most enriched 8-mer sequences (ranked by enrichment) for each polyamide, using MEME [38,39]. MEME was run with following parameters:

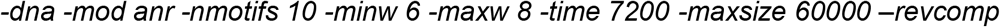

### Specificity and energy landscapes

Specificity and Energy Landscapes (SELs) display high-throughput protein-DNA binding data (DNA–protein interactome or DPI) in the form of concentric rings [40–42]. The organization of data in SEL is detailed in **S3 Fig**. SELs were generated from 8-mer enrichment files using the target sequence for corresponding polyamide **1**(5’-WGWWCW-3’) and **2**(5’-WGGWCW-3’) as seed motif. The software for generating SELs is made available online (https://ansarilab.biochem.wisc.edu/computation.html).

### Genomescapes: scoring *in vivo* bound sites with *in vitro* data

Genomescapes are generated by assigning *in vitro* CSI intensities (enrichment values) to genomic regions. To generate CSI Genomescapes a sliding k-mer window was used to score genomic regions and then plotted as a bar plot [40,42].

### Summation of sites model

Summation of sites (SOS) model was used to predict DNA binding of polyamides in the human genome, hg19 [23,24,43]. The SOS score was obtained by summing (or averaging) all k-mer *in vitro* binding intensities (enrichment) obtained using a sliding k-mer window across a genomic region. Data is displayed using genomic regions of 420 bp for SOS [23]. For SOS predicted genomic loci, the whole human genome (hg19) was divided into 420 bp fragments with the overlap of half (210 bp). These fragments were then sorted by the predicted binding to polyamide **1** and **2** using SOS model. The top 1000 predicted peaks obtained were used as final predicted peaks for further analysis.

### COSMIC-seq data analysis

Sequencing reads were mapped to the human genome (hg19) with Bowtie (best -m 1) to yield unique alignments. Bound regions/peaks were identified with SPP [24,43]. The data has been deposited in the Gene Expression Omnibus (GEO) database, www.ncbi.nlm.nih.gov/geo (accession no. GSE149367).

### Plotting tag density data, genomescape and SOS as heatmaps

To display data multiple heatmaps are shown using two types of genomic regions: the top 1000 COSMIC peaks, and the top 1000 genomic loci predicted to be the best binder using the SOS model. These regions were scored using COSMIC-seq tag density for AP of a 10 Kbp region surrounding the peak using HOMER annotatePeaks.pl command with arguments *-hist as 25 bp*, SOS scores for 10 Kbp region surrounding the peak, and genomescapes for 1 Kbp [44]. Different coloring scales were used to display heatmaps by using a multiplication factor of 10x for tag density and 100x for genomescapes and SOS scoring.

## RESULTS

### Polyamide Design

COSMIC-seq was performed on two structurally identical hairpin polyamides, differing at a single position (X = CH vs N:) on the second aromatic amino acid ring-pair, **Fig 1A**. A single CH to N: position substitution changes the ring pair from a Py/Py to an Im/Py which invokes a preference from an A•T or T•A to a G•C base pair, respectively, based on previously determined pairing rules, **Fig 1B** [1,2,45]. Py-Im polyamide **1**, designed to target the consensus androgen response element (ARE) half-site 5’-WGWWCW-3’, has been shown to regulate androgen receptor (AR) and glucocorticoid (GR) driven gene expression in cell culture and suppress tumor growth *in vivo* [14,18,46]. Py-Im polyamide **2**, designed to target the estrogen response element (ERE) consensus half site 5’-WGGWCW-3’, was shown to effect estrogen receptor-alpha (ERα)-driven gene expression *in vitro* and *in vivo* [47]. In this study each, hairpin Py-Im polyamide is conjugated at the C-terminus with a psoralen and biotin for enrichment connected via a linker (~36 Å extended) capable of sampling pyrimidine proximal to the polyamide-binding site suitable for 2 + 2 photocycloaddition [23,24]. Because the psoralen moiety crosslinks proximal pyrimidines (T in particular), we anticipate subtle bias in the data, a contextual flanking sequence nuance adjacent to the core binding sites of each polyamide. Py-Im polyamides were synthesized by Boc solid-phase synthesis, cleaved from resin and conjugated to the psoralen-biotin moiety 3 (**S1 Fig**).

### Different sequence specificities conferred by a single position substitution

To comprehensively map *in vitro* binding characteristics of hairpin polyamide-conjugates **1** and **2**, we performed solution-based Cognate Site Identifier (CSI) analysis, **Fig 2** [6,40,41,48]. Sequence specificity data was determined with next generation sequencing (NGS) by solution-based enrichment methods (SELEX-seq) to assess polyamide-DNA binding [49]. These methods provide a comprehensive characterization of polyamide-DNA binding through the sampling of a large sequence space (a dsDNA library bearing all ~10^12^ sequence permutations of a 20-bp site) using affinity purification coupled with massively parallel sequencing [35,41]. This platform allows rapid, quantitative identification of the full spectrum of polyamide binding sites of up to 20 bp in size, correlates well with solution-phase and microarray platforms, and has been used to guide the refinement of general polyamide design principles [9,24,31,40,41]. Py-Im polyamides **1** and **2** were incubated with a duplex oligonucleotide library containing a randomized 20-mer region, and the bound and unbound sequences were separated via affinity purification by streptavidin-coated magnetic beads, **Fig 2A**. Following each round of enrichment, sequences were PCR amplified, purified, multiplexed, and subjected to massively parallel sequencing analysis. Computational analysis was applied to enriched sequences to obtain binding site intensity values corresponding to all 8-mer DNA sequences, see methods [37].

**Fig 2.**
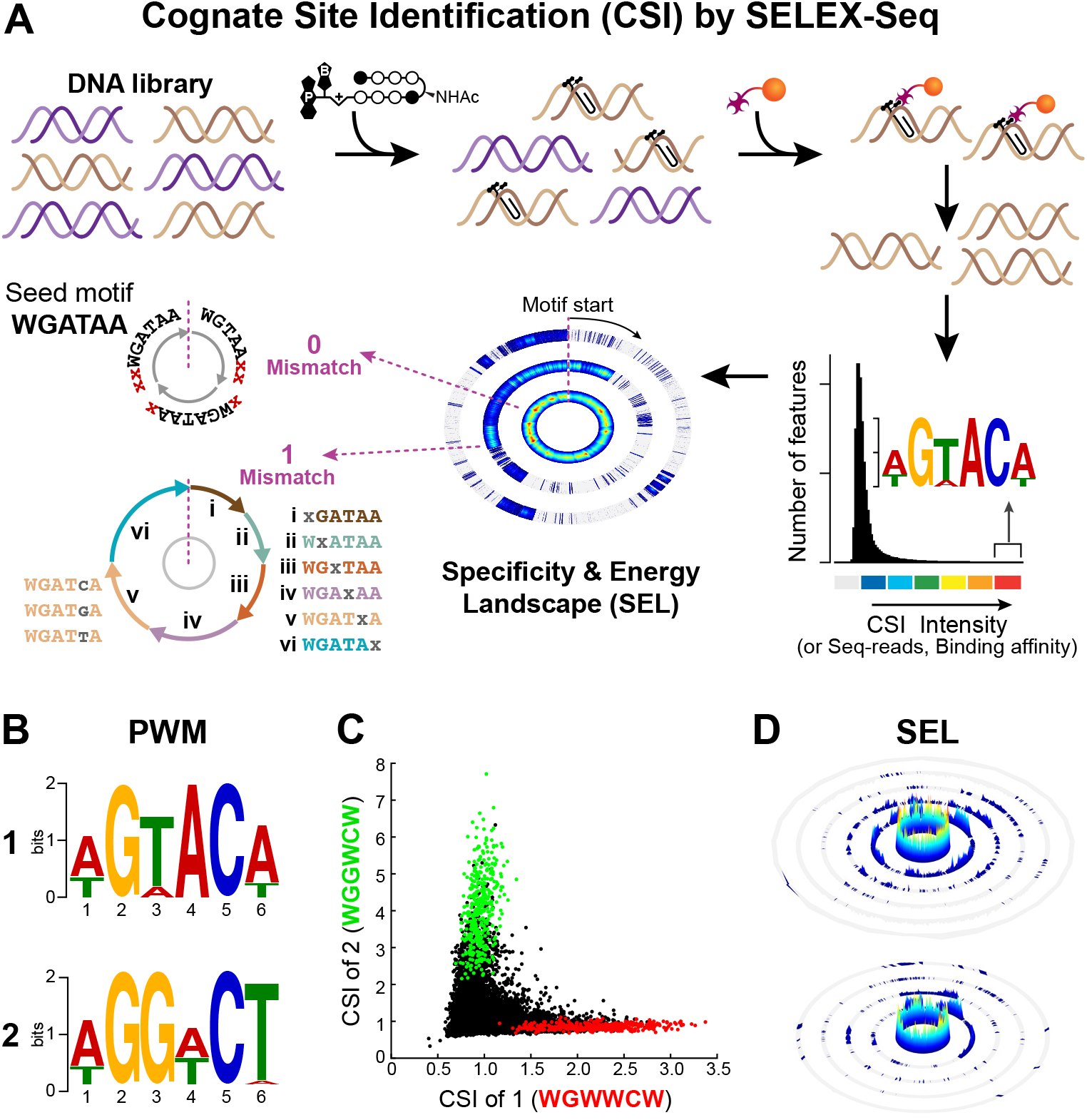
Detection of *in vitro* DNA binding of polyamides 1 and 2 via cognate site identification (CSI) by SELEX-seq. (**A**) Overview of CSI by SELEX-seq workflow. A randomized 20 bp DNA library is incubated with biotinylated polyamide, DNA is enriched by streptavidin-coated magnetic beads, PCR amplified and sequenced by NGS to obtain k-mers (CSI enrichment) representing polyamide-DNA binding. Enrichment is displayed as a histogram plot and high binding sequences are represented as a position weight matrix (PWM) logo. Specificity and energy landscapes (SELs) are created to visualize the full spectrum of DNA binding across all sequence permutations of an 8-mer binding site. (**B**) PWM logos for polyamides **1** (*top*) and **2** (*bottom*). (**C**) Scatterplot comparison of *in vitro* DNA binding for **1** vs **2**. CSI enrichment for 8-mers is plotted for sequences containing 5’-WGWWCW-3’ (*red*) and 5’-WGGWCW-3’ (*green*). (**D**) Comprehensive SELs for **1** (*top*) and **2** (*bottom*) using 5’-WGWWCW-3’ and 5’-WGGWCW-3’ as seed motif, respectively, where W = A or T.

Polyamide-DNA binding motifs for each round of enrichment were identified by position weight matrices (PWMs) using the top 50 enriched 8-mer sequences and displayed as sequence logos, **Fig 2B** and **S2 Fig**, [38,39]. The highest information content for each polyamide is found at a binding site width of six, verifying the binding site size expected when **1** and **2** are bound in a fully ring-paired, hairpin configuration. The motifs generated are indicative of polyamide-DNA binding consistent with the Py-Im pairing rules for both **1** and **2**, targeting the sequences 5′-WGWWCW-3′ and 5′WGGWCW-3′, respectively. A clear difference in the sequence preference at the third position, corresponding to the CH to N: position substitution of the second ring pair (Py/Py vs Im/Py), was detected. Additionally, both polyamides show subtle differences in binding preference at the fourth and sixth positions revealing sensitivity of binding energetics to changes in sequence context. Scatter plot comparison analysis of all enriched 8-mer sequences indicates a preference for the consensus motif, polyamide **1** prefers WGWWCW over WGGWCW, whereas polyamide **2** prefers WGGWCW over WGWWCW, **Fig 2C** and **S3 Fig**. These results demonstrate that a single position (CH to N:) modification of the aromatic amino acid ring of the polyamide core structure imparts a significant change in the global *in vitro* DNA sequence preferences and confirms that the C-terminus modification does not have significant impact on the specificity of the hairpin polyamides. While based on the pairing rules these results may seem obvious, this experiment was important to confirm that polyamide-conjugates **1** and **2** retain preference for cognate sequences.

While PWM-based motifs summarize sequence preferences of DNA-binding molecules, they compress related sequences into a consensus motif, masking the impact of flanking sequences and local microstructure as well as underestimating the affinity spectrum of cognate sites contained within a given DNA-polyamide interactome (DPI). Sequence specificity landscapes (SSLs) can optimize the cognate site motif(s) and thereby uncover major binding motifs to visualize the effects of flanking sequences [40]. To better visualize the full spectrum of DNA binding and compare the individual interactomes of each polyamide, we developed specificity and energy landscapes (SELs) for **1** and **2**, **Fig 2D** [40–42]. SELs present the enriched binding sequences as concentric rings, organized by a “seed motif” in the zero-mismatch ring (central ring) having an exact match for the seed motif, **Fig S4**. The PWM-based motif is used as a seed and the entire DPI displayed in concentric rings as they deviate from the seed motif. Each consecutive ring represents 0, 1, 2, n.., mismatches from the seed motif. SELs plotted for the complete set of enrichment data using a 6-mer seed motif, WGWWCW (**1**) and WGGWCW (**2**), show a clear preference of both polyamides for 6-mer seed motifs (central ring). The dramatic drop-off in affinity for sequences that deviate from the preferred 8-mer site (outer rings), underscores the exquisite sequence specificity of hairpin polyamides, **Fig 2D** and **S5 Fig** [40–42].

It is important to note that low-affinity sequences, that are sequentially depleted in SELEX-based approaches, are critical to develop accurate binding site models across the genome [23,37,40,42]. Indeed, we observed both a concentration and selection effect, with each sequential round of SELEX steadily enriching high-affinity sites with a concomitant decrease in correlation with genome-wide binding profiles (**S1 File**). For these reasons, DNA-polyamide interactome from round 1 (with no successive rounds of SELEX) with final concentration of polyamides at 50 nM was used for further analysis.

## COSMIC-seq to map genome-wide binding profiles

We utilized COSMIC-seq to map the genome-wide binding targets of **1** and **2** in LNCaP nuclei to determine if Py-Im polyamides could maintain their preferred differential binding specificity in a biochemically active complex chromatin environment, **Fig 3A** [24]. Isolated nuclei retain native chromatin states and are widely used to examine chromatin structure and accessibility [50–52]. Briefly, isolated nuclei from human LNCaP cells were treated in biological duplicate with **1** or **2** (0.4 μM and 4 μM) at 4 °C for 1 h and cross-linked to DNA by 365 nm UV irradiation. DNA was sheared by sonication, polyamide-DNA complexes were captured by streptavidin-coated magnetic beads, cross-links were reversed, and enriched DNA was sequenced. COSMIC-seq reads were mapped using the Bowtie algorithm and bound peaks were identified using the standard peak-calling algorithms endorsed by the ENCODE consortium, see Methods [24,43,53].

**Fig 3.**
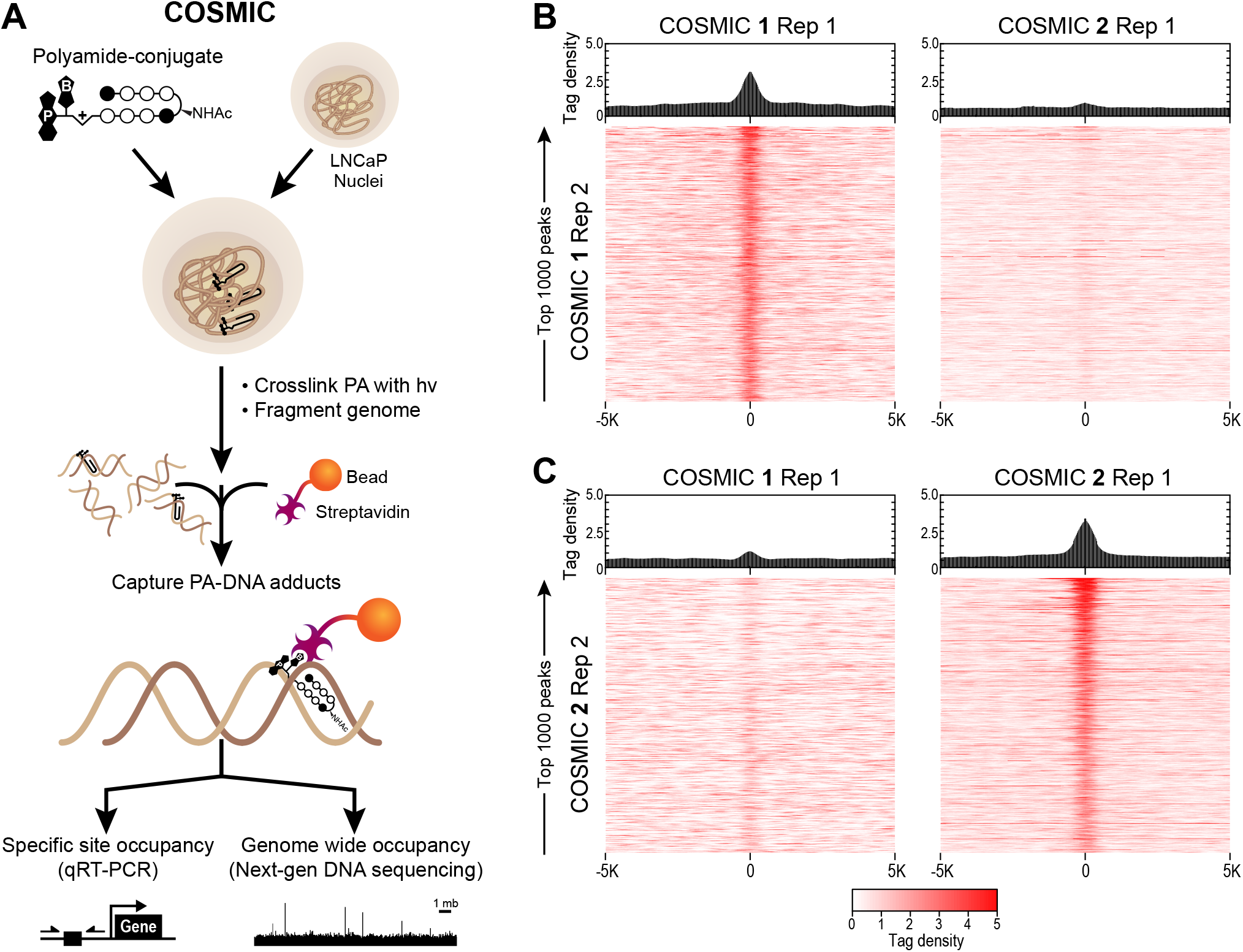
Genome-wide DNA binding of polyamides 1 and 2 by COSMIC-seq. (**A**) Overview of COSMIC-seq in LNCaP cells, nuclei are treated with polyamides **1** and **2** and cross-linked to DNA with UV irradiation (365 nm). Cross-linked genomic DNA is enriched and analyzed by NGS. (**B**, **C**) Heat maps reveal selective enrichment of polyamides **1** and **2**. Tag density of each polyamide is shown for the top 1,000 loci for **1**(**B**) and **2**(**C**). Data is displayed as sequence read tag density heatmaps (*bottom*) and averaged bar plots (*top*) for the top 1000 predicted peaks are mapped on a 10 Kbp window.

Sequence/Read tag density of the top 1000 identified peaks across replicates of polyamides was compared over a 10 Kbp region centered at bound COSMIC-seq loci, and shown as a heat map, **Fig 3B** and **3C**. Both polyamides **1** and **2** show a strong correlation between bound peaks of replicates while a consistent non-correlation is observed when comparing the top 1000 identified peaks of polyamide **1** to those of polyamide **2**. A high COSMIC enrichment signal for sites bound by polyamide **1** is observed for replicate treatments (**Fig 3B**, *left*), however, no enrichment is observed when compared to polyamide 2 (**Fig 3B**, *right*). A similar negligible enrichment overlap is observed for sites bound by polyamide 2 (**Fig 3C**, *left*), while a clear enrichment between polyamide **2** replicate treatments is identified (**Fig 3C**, *right*). Consistent with the *in vitro* binding analysis (**Fig 2C**) polyamide **1** and **2** show poor overlap at bound genomic loci in LNCaP nuclei. These results indicate that differential polyamide sequence specificity is maintained in the context of the native chromatinized genome.

## Genomic occupancy correlates with binding predictions based on summation of sites (SOS) model

An overview of the COSMIC (genomic occupancy) and CSI (*in vitro* specificity) pipelines and the comparative analysis of the two datasets is shown in **Fig 4A** (see also Methods). CSI genomescapes are generated by assigning binding probability scores across the entire human genome. Unlike standard PWM-based genome annotation approaches, CSI-Genomescapes take into account potential moderate-to-low affinity cognate sites [24,40]. Binding intensity is assigned to every 8-bp sequence in the genome based on the CSI data (*in vitro*), and compared to the top 1,000 COSMIC peaks (*in nuclei*) over a 1 Kbp region (**Fig 4A**, *right*). While the average predicted binding is higher at identified COSMIC loci for polyamides, signal resolution is low when attempting to compare *in vitro* binding at individual COSMIC peaks, **S6 Fig**. A summation of sites (SOS) scoring model is a more robust method for predicting *in vivo* binding by considering clusters of potential binding sites [24]. Recent studies suggest that genome-wide binding events for natural DNA-binding transcription factors as well as engineered small molecules occur at genomic loci bearing clusters of high-affinity and multiple moderate and weak-affinity binding sites [23]. The CSI *in vitro* binding data of **1** vs **2** clearly illustrate a differential preference for flanking sites and tolerated deviations from seed motifs. These differences could potentially affect the genomic-binding and are known to influence genome-wide distributions of architecturally different polyamides (hairpin versus linear structures). SOS scoring was used to predict binding potential for **1** and **2** across the human genome [23]. By comparing COSMIC signals with predicted binding based on CSI-derived genomescapes, we observed that the sum of all *in vitro* determined binding intensities (Z-scores), tiled across an ∼420-bp window, most reliably predicted polyamide occupancy at distinct genomic loci (**Fig 4B**, **S7 Fig**, and **S2 File**). There is a strong correlation between the top 1000 COSMIC peaks (*in nuclei*) and SOS signals (*in vitro*) for both polyamide **1** and **2**. Notably, there is no correlation observed when comparing *in vitro* and COSMIC data between polyamide **1** and **2**.

**Fig 4.**
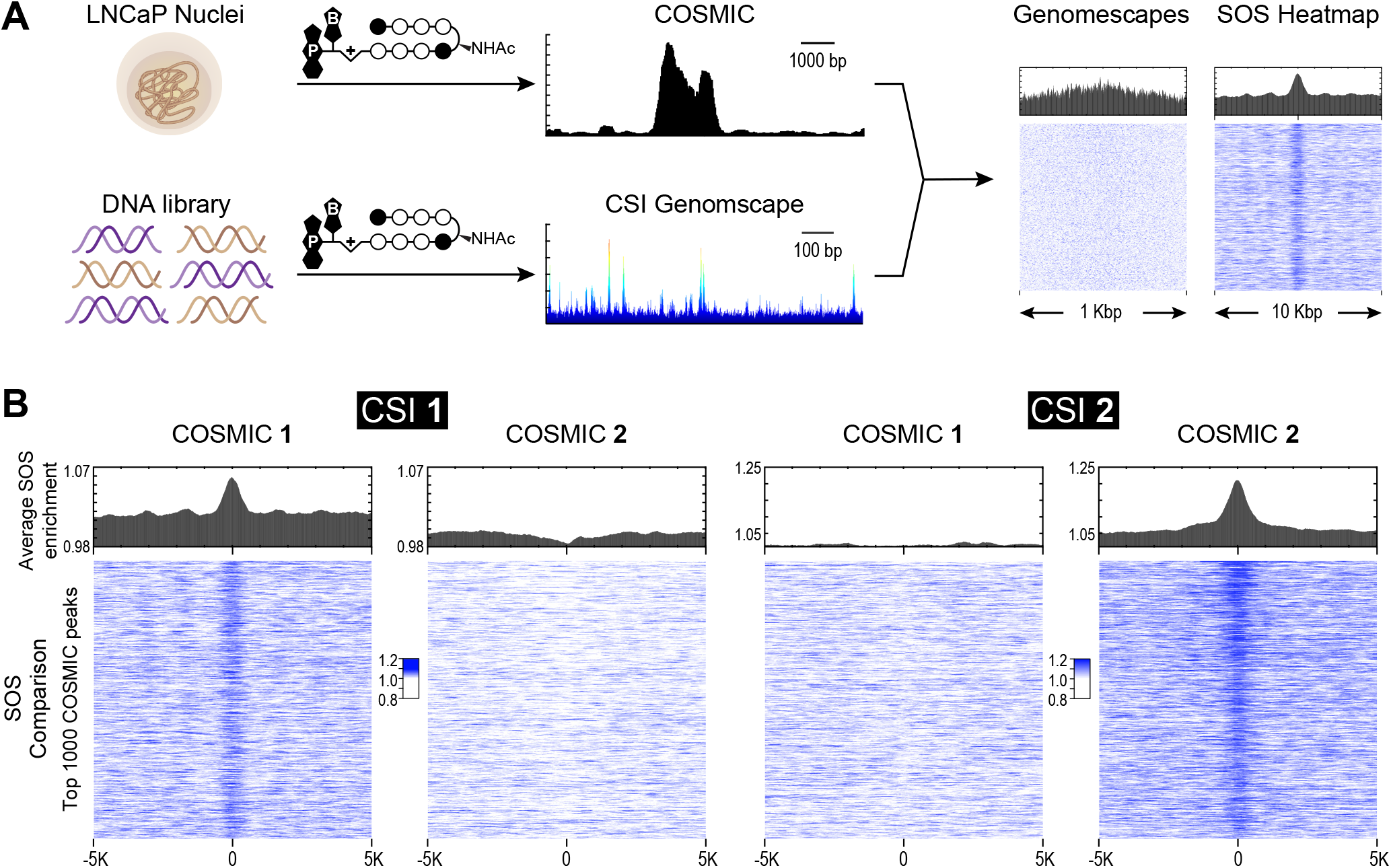
Polyamide binding across the genome correlates to *in vitro* binding predictions. (**A**) COSMIC-scape analysis generates CSI genomescapes and SOS heatmaps using the CSI-SELEX and COSMIC-seq data. (**B**) Data is displayed as averaged bar plots (*top*) and SOS heatmaps (*bottom*) for top 1000 COSMIC peaks mapped on a 10 Kbp region. SOS heatmaps demonstrate that *in vitro* binding of polyamides **1**and **2** is predicted at COSMIC-seq loci while there is no correlation observed between polyamides.

To access binding differences at individual loci, we selected loci identified by COSMIC on chromosomes 2 and 19 (chr2 and chr19), which were predicted to bind **1** and **2**, respectively, **Fig 5**. By comparing CSI 8-mer enrichment profiles and SOS profiles (**Fig 5A** and **5B**) to the COSMIC tag density (**Fig 5C** and **5D**) for both polyamides, it is evident that polyamides **1** and **2** have distinct, non-overlapping genomic binding preferences. Consistent with sequence-specific binding, polyamide **1** is not found at the CSI-predicted loci on chr19 for polyamide **2** (**Fig 5C**), and vice versa, **2** is not found at the chr2 predicted loci for **1** (**Fig 5D**). A similar example of a relationship between loci on chromosome 10 is displayed in **S8 Fig**. These studies clearly demonstrate a non-correlation between the COSMIC and CSI of **1** vs **2**, and a distinct correlation between *in nuclei* (COSMIC) and *in vitro* (CSI) DNA binding properties.

**Fig 5.**
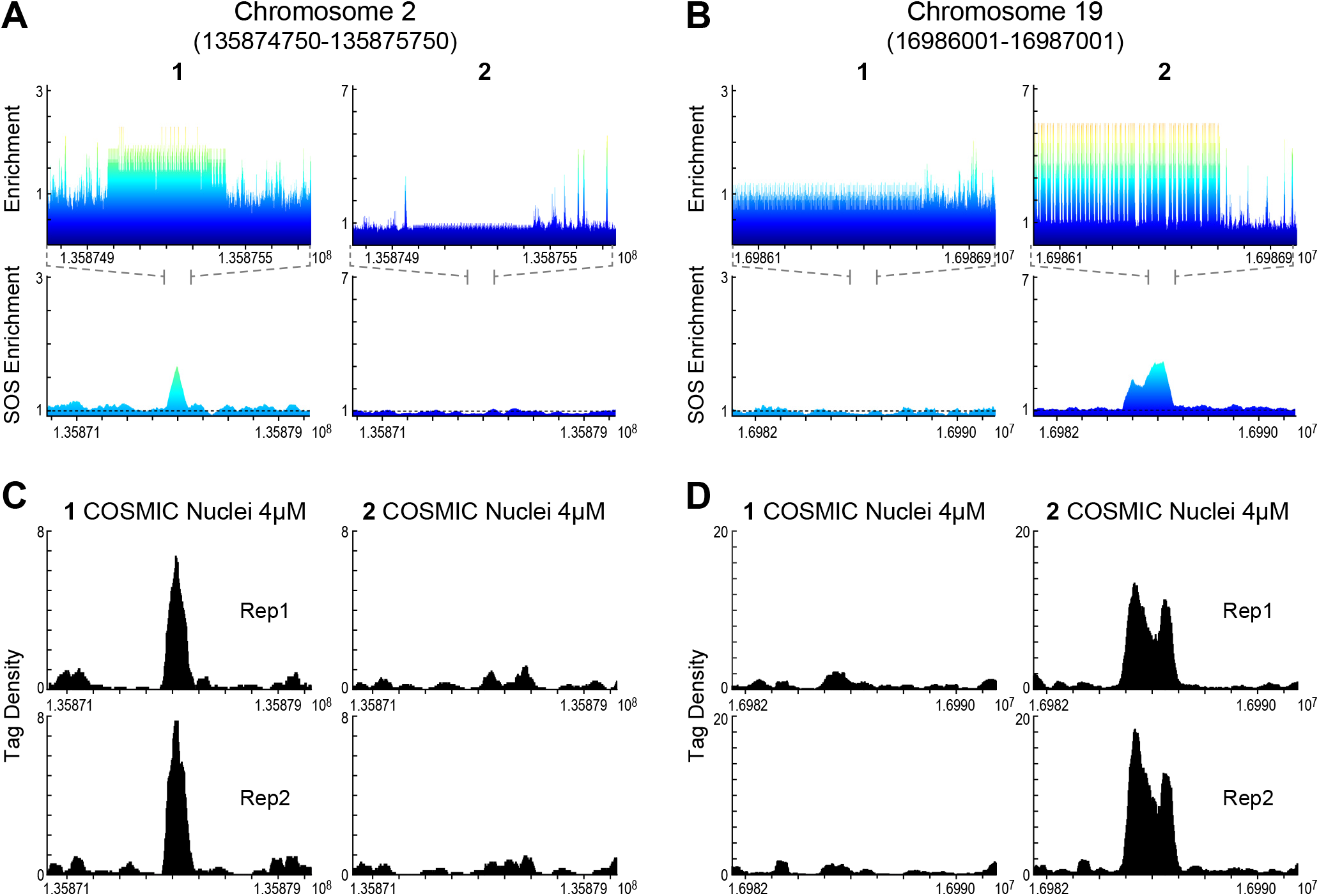
Polyamides 1 and 2 have distinct, non-overlapping genomic binding preferences. (**A**, **B**) Genomescapes (*top*) displaying a 1 Kbp region and SOS enrichment plots (*bottom*) displaying a 10 Kbp region for polyamides **1** and **2** at genomic loci of chr2 and chr19, respectively. (**C**, **D**) COSMIC tag density data of replicates of **1**and **2** for a 10 Kbp region at same loci.

## DISCUSSION

We determined the genome-wide binding events of Py-Im polyamides of similar hairpin architecture that differ at a single position (CH vs N:) on the second aromatic amino acid ring-pair. This single position substitution changes the ring pair from a Py/Py to an Im/Py which alters the binding preference from an A•T or T•A to a G•C base pair, respectively. Comprehensive *in vitro* binding analyses confirm the preferential binding of polyamide **1** to WGWWCW and polyamide **2** to WGGWCW [7]. Both polyamides exhibited high selectivity for their cognate motifs, while exhibiting considerably lower binding affinity for the motif of the other hairpin polyamide. These observations are indicative of polyamide-DNA binding consistent with the established Py-Im pairing rules for both **1** and **2** [54,55]. When comparing binding events within LNCaP nuclei, we see correlation among replicates using the SOS model. Additionally, consistent with the *in vitro* binding analysis, hairpin **1** and **2** show poor overlap in bound genomic loci indicating that innate sequence preferences of these structurally similar hairpin polyamides are maintained in the context of the chromatinized nuclear genome. These results demonstrate that a single position (CH to N:) modification of the aromatic amino acid ring of the polyamide 8-ring structure imparts a significant change in binding preference that is maintained within cellular nuclei.

We chose to examine genome-wide binding profiles in intact cell nuclei not only because they present compacted genomic DNA in a chromatinized context but they also circumvent the complexity of cellular uptake of high molecular weight polyamide conjugates. Similarly, state-of-the-art genomic chromatin structure and accessibility studies in a wide range of cell- and tissue-types primarily rely on isolated nuclei because they have been demonstrated to accurately capture native chromatin states in living cells [50–52].

Py-Im polyamides have been shown to modulate oncogenic transcription factor signalling and reduce their binding occupancy at select loci in ChIP experiments [14]. A recent study demonstrated that a polyamide targeting dihydrotestosterone (DHT) inducible AR-DNA binding was able to repress 30% of DHT inducible binding events [56]. Importantly, motif analysis of the repressed AR peaks demonstrated that the differential effects on AR-DNA binding events *in vivo* reflects the DNA target sequence binding preference of the hairpin polyamide *in vitro*. Consistent with previous work, we observe a strong correlation between genome-wide SOS scoring using polyamide *in vitro* binding data with the corresponding COSMIC genomic binding data [24].

The ability to modify a hairpin Py-Im polyamide while maintaining specificity in a chromatinized environment is an encouraging finding for application as synthetic transcription factors (Syn-TFs). Syn-TFs comprise a modular DNA-binding domain directly fused to or designed to recruit a regulatory domain capable of modulating gene expression when localized to genomic regulatory elements [57,58]. Protein-based artificial TFs have been developed based on zinc finger proteins (ZFPs), transcription activator-like effectors (TALEs), and the clustered regularly interspaced short palindromic repeat (CRISPR)-associated (Cas) system [57,58]. However, these methods may be limited by the lack of an efficient delivery mechanism, *in vivo* bioavailability and unknown immunogenic factors [59–61]. The use of small molecule syn-TFs is an attractive non-protein alternative to regulating transcription allowing for more finely tuned control of dosage and timing without the need for complex genomic integration. [43,62–66]. Small molecule solutions have the potential to be used as molecular tools to dissect endogenous gene networks, epigenetic landscapes and as therapeutics to modulate adherent gene expression.

## Acknowledgements

We thank Laura Vanderploeg for help with the artwork and Mackenzie C. Spurgat for preliminary CSI experiments. Sequencing was performed at the Millard and Muriel Jacobs Genetics and Genomics Laboratory at California Institute of Technology.

## Supporting information

**S1 Fig. Chemical Synthesis and characterization of Py-Im polyamide conjugates. (A)** Solid phase synthetic scheme for the synthesis of Py-Im polyamides **1A** and **2A**, **a)** Boc-Py-OBt, *i*-Pr₂NEt, DMF, μW (80 °C, 3 h); **b)** 80:1:19 TFA:triethylsilane:CH_2_Cl_2_, 5 min, RT; **c)** Boc-Py-OH, PyBOP *i*-Pr₂NEt, DMF, μW (60 °C, 5 min); **d)** 9:2:1 DMF:Ac_2_O:*i*-Pr₂NEt, 30 min, RT; **e)** repeat (1x) steps b - d; **f)** 80:1:19 TFA:triethylsilane:CH_2_Cl_2_, 5 min, RT; **g)** Boc-Im-OH, PyBOP, *i*-Pr₂NEt, DMF, μW (60 °C, 5 min); **h)** 9:2:1 DMF:Ac_2_O:*i*-Pr₂NEt, 30 min, RT; **i)** 80:1:19 TFA:triethylsilane:CH_2_Cl_2_, 25 min, RT; **j)** Fmoc-D-Dab(Boc)-OH, PyBOP, *i*-Pr₂Net, DMF, μW (60 °C, 25 min); **k)** 9:2:1 DMF:Ac_2_O:*i*-Pr₂NEt, 30 min, RT; **l)** repeat (2x) steps b - d; **m)** 80:1:19 TFA:triethylsilane:CH_2_Cl_2_, 5 min, RT; **n) 5**, PyBOP, *i*-Pr₂NEt, DMF, μW (60 °C, 5 min); **o)** 20% piperidine, DMF, 30 min, RT; **p)** 9:2:1, DMF:Ac_2_O:*i*-Pr₂NEt 30 min, RT; **q)** neat 3,3′-Diamino-N-methyldipropylamine, μW (60 °C, 10 min); **(B)** Synthesis of the psoralen–biotin-acid moiety **3**, **a)** Fmoc-PEG_2_-OH, *i*-Pr₂NEt, DCM; **b)** 20% piperidine, DMF; **c)** Biotin-Lys(Fmoc)-OH, HATU, HOAt, *i*-Pr₂NEt, 3:1 DMSO:DMF; **d)** 20% piperidine, DMF; **e)** Fmoc-PEG_2_-OH, HATU, HOAt, *i-*Pr₂NEt, DMF; **f)** 20% piperidine, DMF; **g)** SPB (NHS-psoralen), *i*-Pr₂NEt, DMF; **h)** 95% TFA, 2.5% H_2_O, 2.5% *i*-Pr3SiH; and **(C)** peptide coupling of Py-Im polyamides **1A** and **2A** with **3**; **(C)** Analytical HPLC traces of **1**, **2**, and **3**; **(D)** Characterization of compounds by MALDI-TOF.

**S2 Fig. Position weight matrix (PWM) motif logo representation obtained from the top 50 sequences via CSI by SELEX-seq for polyamides conjugates.** PWMs for three replicates of **1** (*left*) and **2** (*right*) at two concentrations (5 nM and 50 nM) and three enrichment rounds (1, 2 and 3). The PWMs are derived using MEME software with the corresponding e-value indicated.

**S3 Fig. Specificity and energy landscapes (SELs) display the comprehensive binding preferences of polyamides based on a seed motif.** The height of each peak corresponds to CSI enrichment for a given sequence. (**A**) SEL for **1** with seed motif WGWWCW (where W = A or T). (**B**) Top view of the SEL in **A**. (**C**) SEL for **2** with seed motif WGGWCW (where W = A or T). (**D**) Top view of the SEL in **C**. (**E**) SELs consists of concentric rings with sequences in the 0 mismatch ring (central ring) having an exact match to the seed motif. Moving outwards, the 1 mismatch ring contains all sequences that differ from the seed motif at any one position (or a Hamming distance of one). In each ring, sequences are arranged clockwise by position of the mismatch, then alphabetically by the sequence. The 1 mismatch ring begins with mismatches at the first position of the motif and ends with mismatches at the last position of the motif.

**S4 Fig. Specificity and energy landscapes (SEL) representation for all k-mer binding enrichment obtained via CSI by SELEX-seq for polyamide conjugates.** SELs for three replicates of **1** (*left*) and **2** (*right*) at two concentrations (5 nM and 50 nM) and three enrichment rounds (1, 2 and 3).

**S5 Fig. Scatter plots and correlation coefficients denoting replicability of CSI replicates for polyamides conjugates.** Scatter plots for CSI enrichment of **1 (A)** and **2 (B)** at two concentrations (5 nM and 50 nM) and three enrichment rounds (1, 2 and 3).

**S6 Fig. COSMIC loci compared to CSI genomescapes.** Data is displayed as averaged bar plots (*top*) and heatmap of genomescapes (*bottom*) for top 1000 COSMIC peaks mapped on a 1 Kbp region. CSI data from enrichment round 1 at 50 nM for **1** (*left*) and **2** (*right*) was used for genomescape generation.

**S7 Fig. COSMIC-seq tag density data plotted for the top 1000 SOS predicted sites shows COSMIC binding is found at the predicted genomic sites.** Heatmaps with tag density for COSMIC replicates of **1**(**A**) and **2**(**B**) are mapped for the top 1000 SOS predicted genomic peaks using a 10 Kbp window. CSI data from enrichment round 1 at 50 nM for **1** (*left*) and **2** (*right*) was used for SOS prediction.

**S8 Fig. Polyamides 1 and 2 have distinct, non-overlapping genomic binding preferences.** (**A**) Genomescapes (*top*) displaying a 1 Kbp region and SOS enrichment plots (*bottom*) displaying a 10 Kbp region for polyamides **1** and **2** at genomic loci of chr10. (**B**) COSMIC tag density data of replicates of **1** and **2** for a 10 Kbp region at same loci.

**S1 File. SOS and genomescapes**. Heatmaps for SOS and genomescape data for enrichment round 1 at 50 nM for polyamide **1** and **2**. Heatmaps are plotted for the top 1000 COSMIC peaks of COSMIC replicates of polyamides **1** and **2** on a 10 Kbp window for SOS and 1 Kbp for genomescapes.

**S2 File. COSMIC-seq tag density data.** Tag density heatmaps for polyamide **1** and **2** replicates are mapped for the top 1000 SOS predicted genomic peaks using a 10 Kbp window. CSI data from enrichment round 1 at 50 nM for **1** and **2** was used for SOS prediction.

