## Supplemental Figures 1-8 for "Single position substitution of hairpin pyrrole-imidazole polyamides imparts distinct DNA-binding profiles across the human genome"

**A**

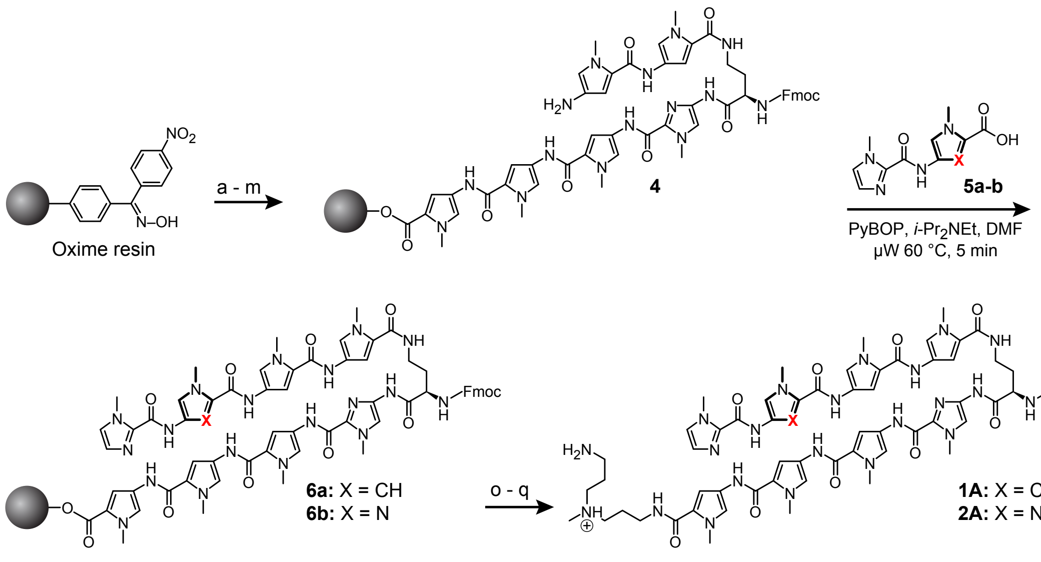
Oxime resin
a - m
4
5a-b
PyBOP, i-Pr2NEt, DMF
μW 60 °C, 5 min
6a: X = CH
6b: X = N
o - q
1A: X = CH
2A: X = N

**B**

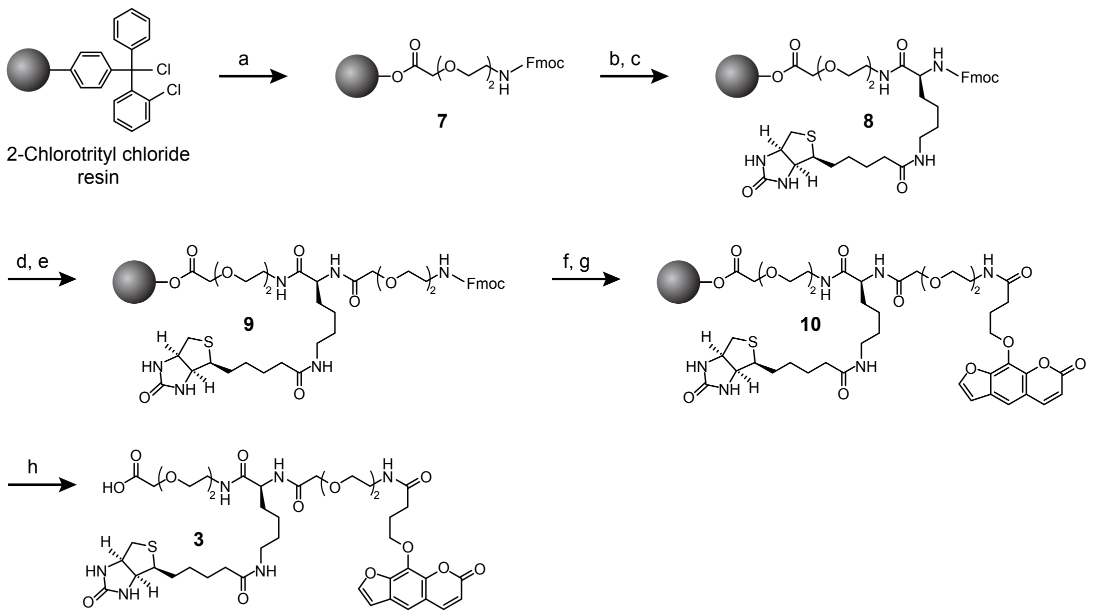
2-Chlorotrityl chloride
resin
a
7
b, c
8
d, e
9
f, g
10
h
3

**C**

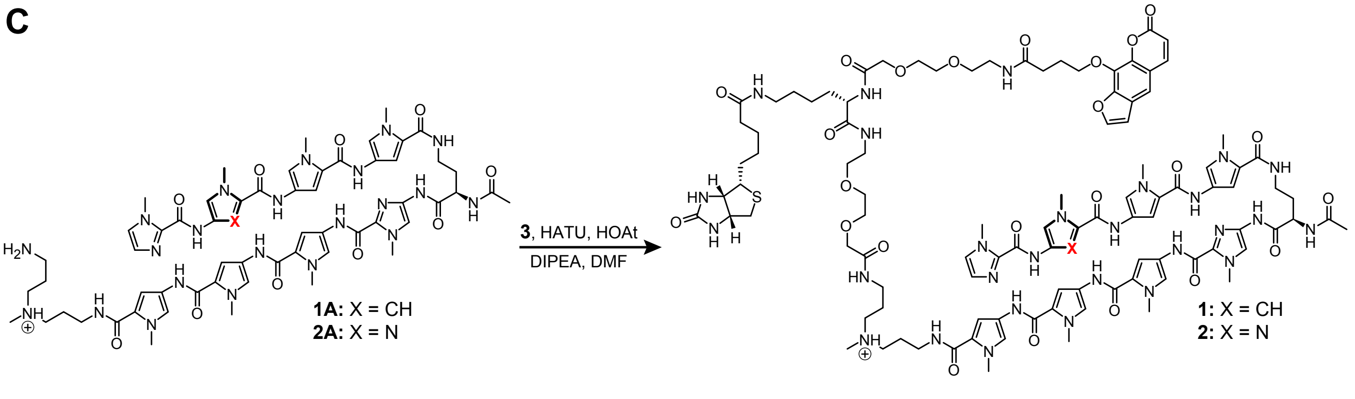
1A: X = CH
2A: X = N
3, HATU, HOAt
DIPEA, DMF
1: X = CH
2: X = N

**D**

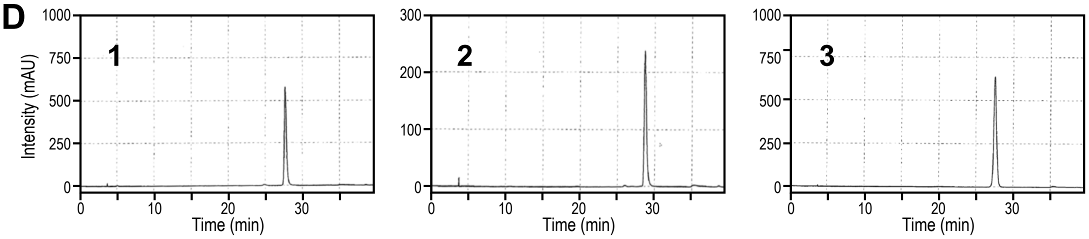


**E**

| Compound | Chemical Formula [M+H]$^+$ | Calculated Mass [M+H]$^+$ | Observed Mass [M+H]$^+$ |
|---|---|---|---|
| **1A** | $C_{59}H_{75}N_{22}O_{10}$ | 1251.60 | 1251.78 |
| **2A** | $C_{58}H_{74}N_{23}O_{10}$ | 1252.60 | 1252.75 |
| **1** | $C_{102}H_{133}N_{28}O_{24}S$ | 2165.98 | 2166.26 |
| **2** | $C_{101}H_{132}N_{29}O_{24}S$ | 2166.97 | 2167.23 |
| **3** | $C_{43}H_{61}N_{6}O_{15}S$ | 933.39 | 933.36 |

Supp. Figure 2

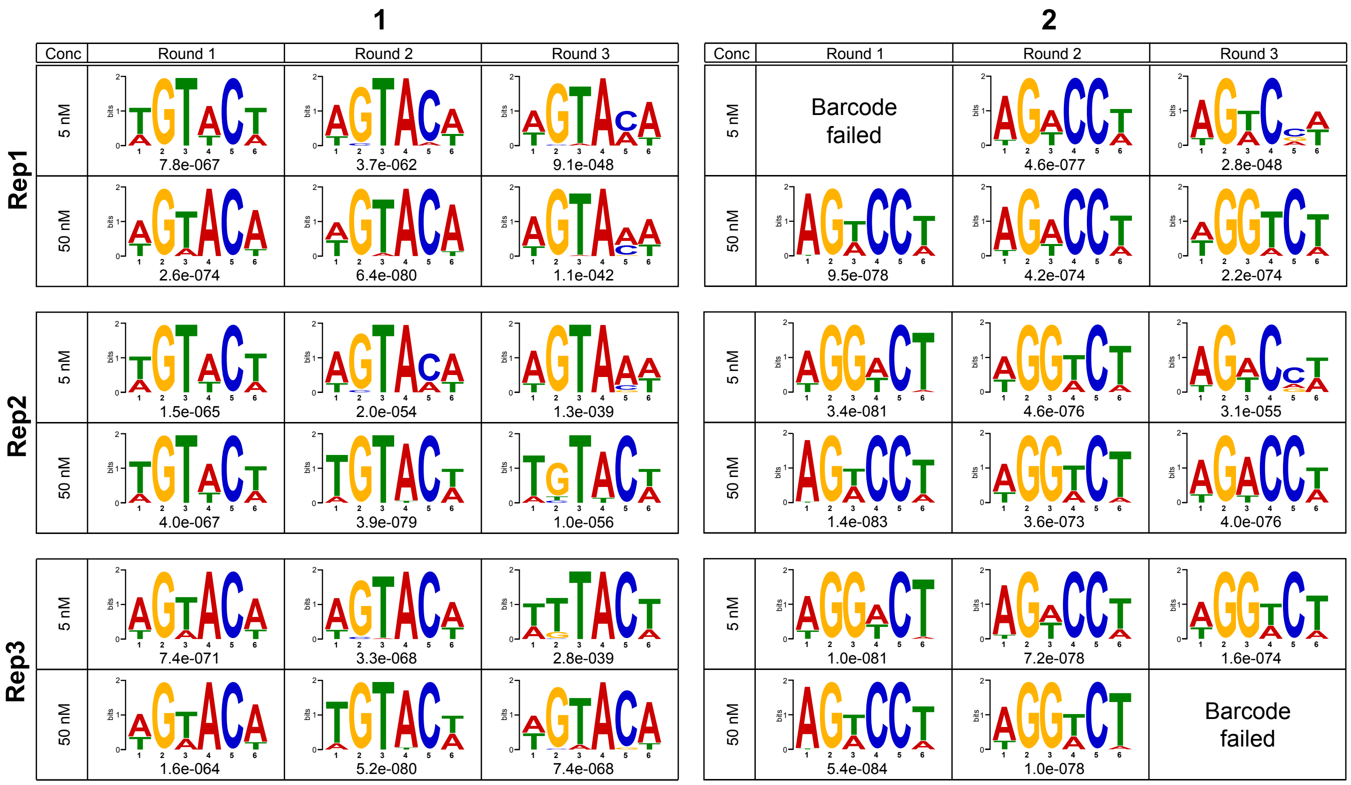

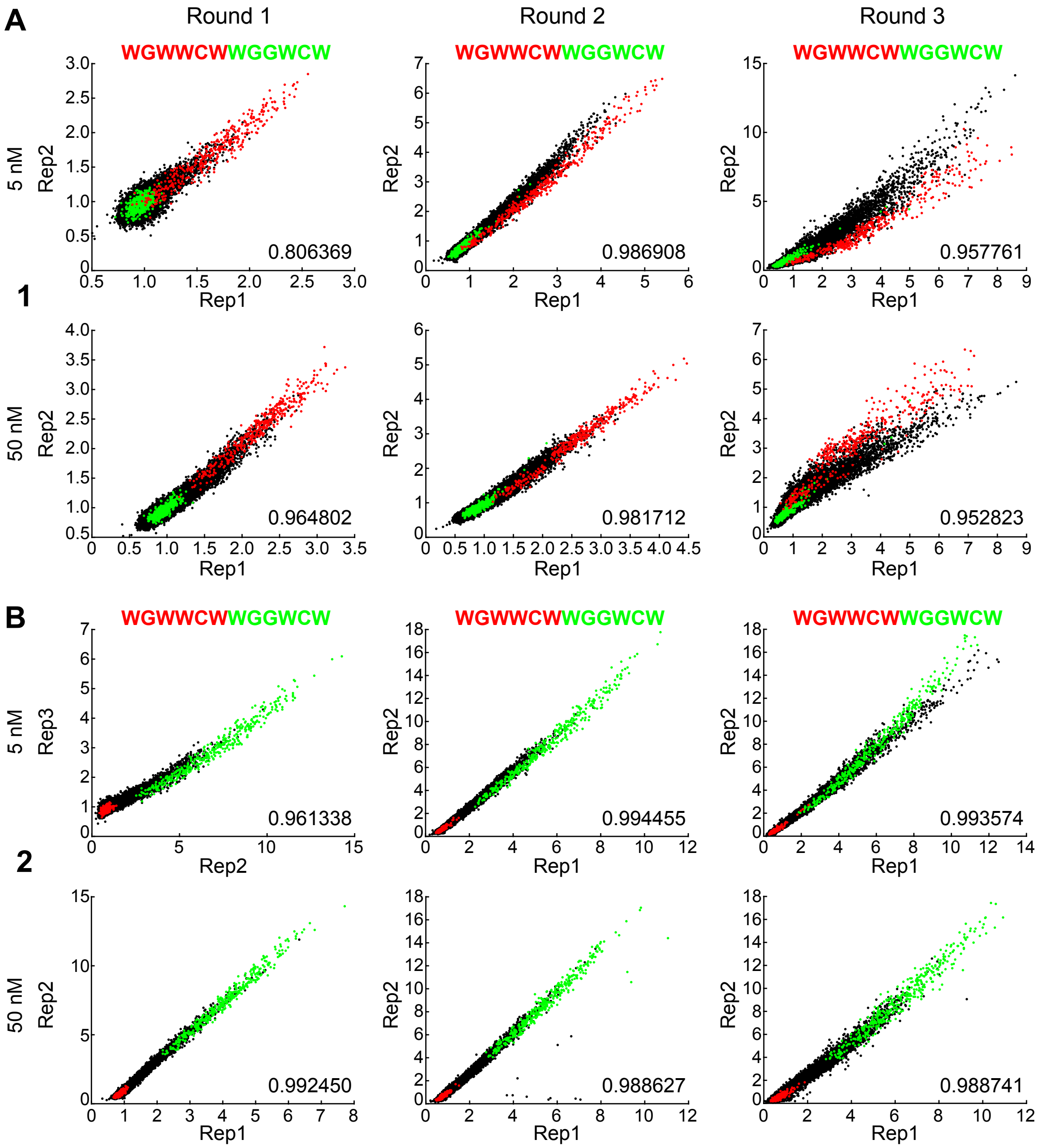
A
B
1
2
Round 1
Round 2
Round 3
WGWWCWWGGWCW
5 nM
50 nM
Rep1
Rep2
Rep3
0.806369
0.986908
0.957761
0.964802
0.981712
0.952823
0.961338
0.994455
0.993574
0.992450
0.988627
0.988741

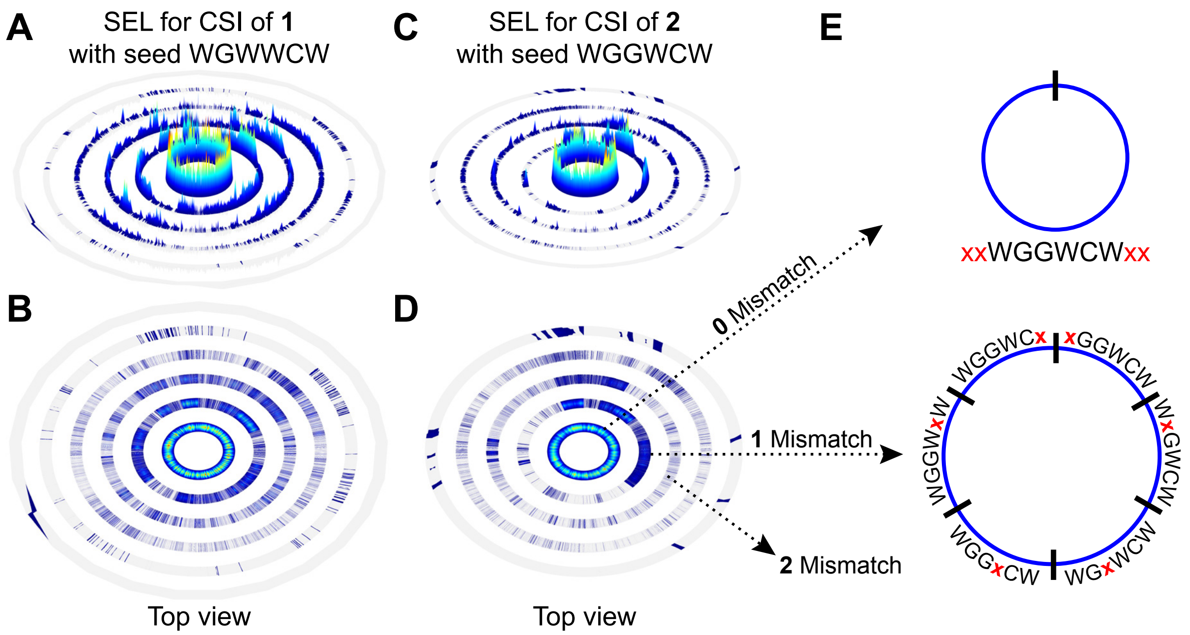
A
SEL for CSI of 1
with seed WGWWCW
C
SEL for CSI of 2
with seed WGGWCW
E
B
D
Top view
Top view
0 Mismatch
1 Mismatch
2 Mismatch
xxWGGWCWxx
WGGWCx
xGGWCW
WxGWCW
WGxWCW
WGGxCW
WGGWxW

### Supp. Figure 5

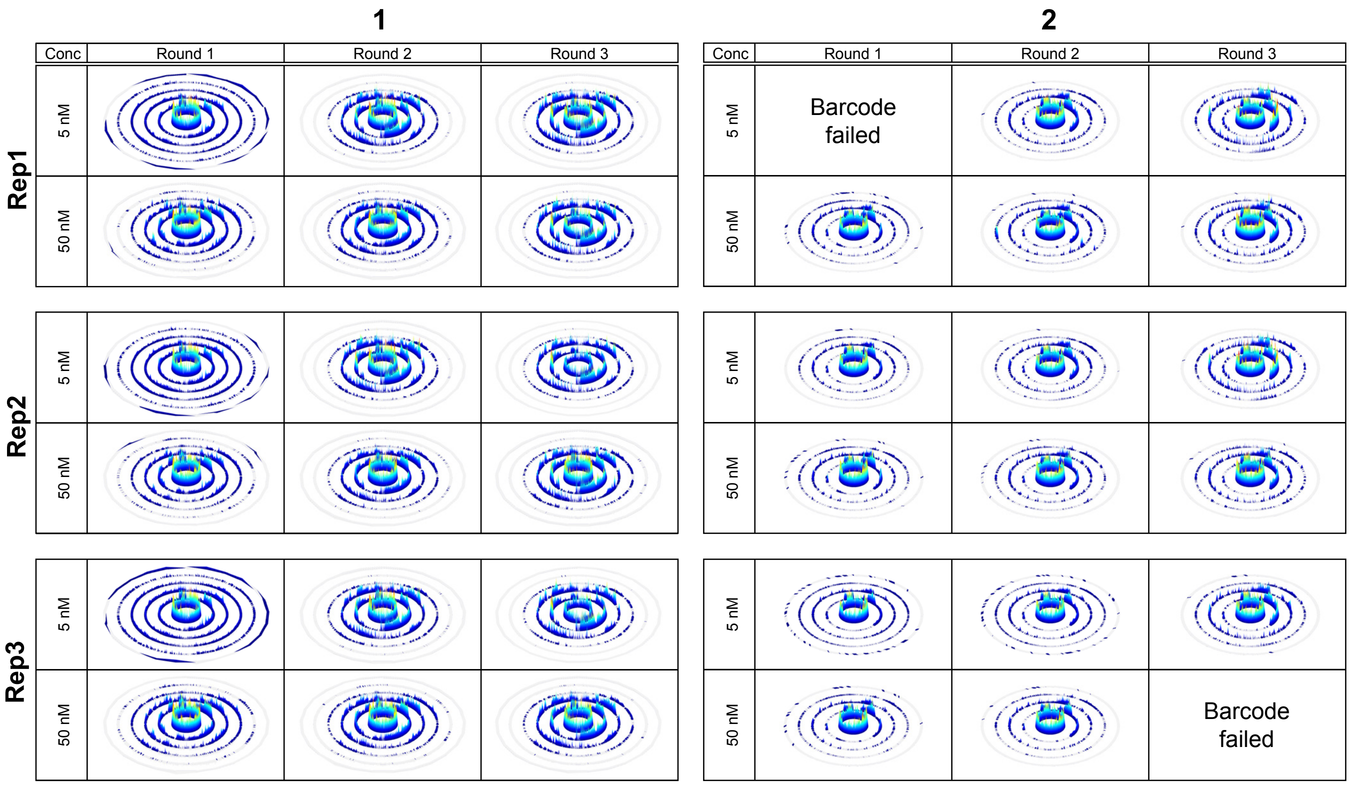

Supp. Figure 6

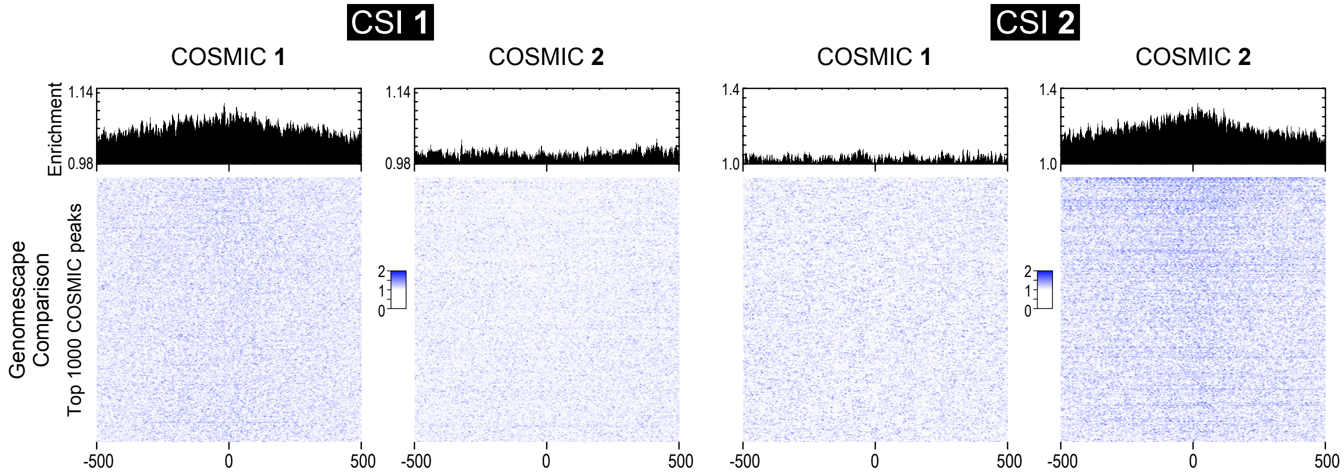

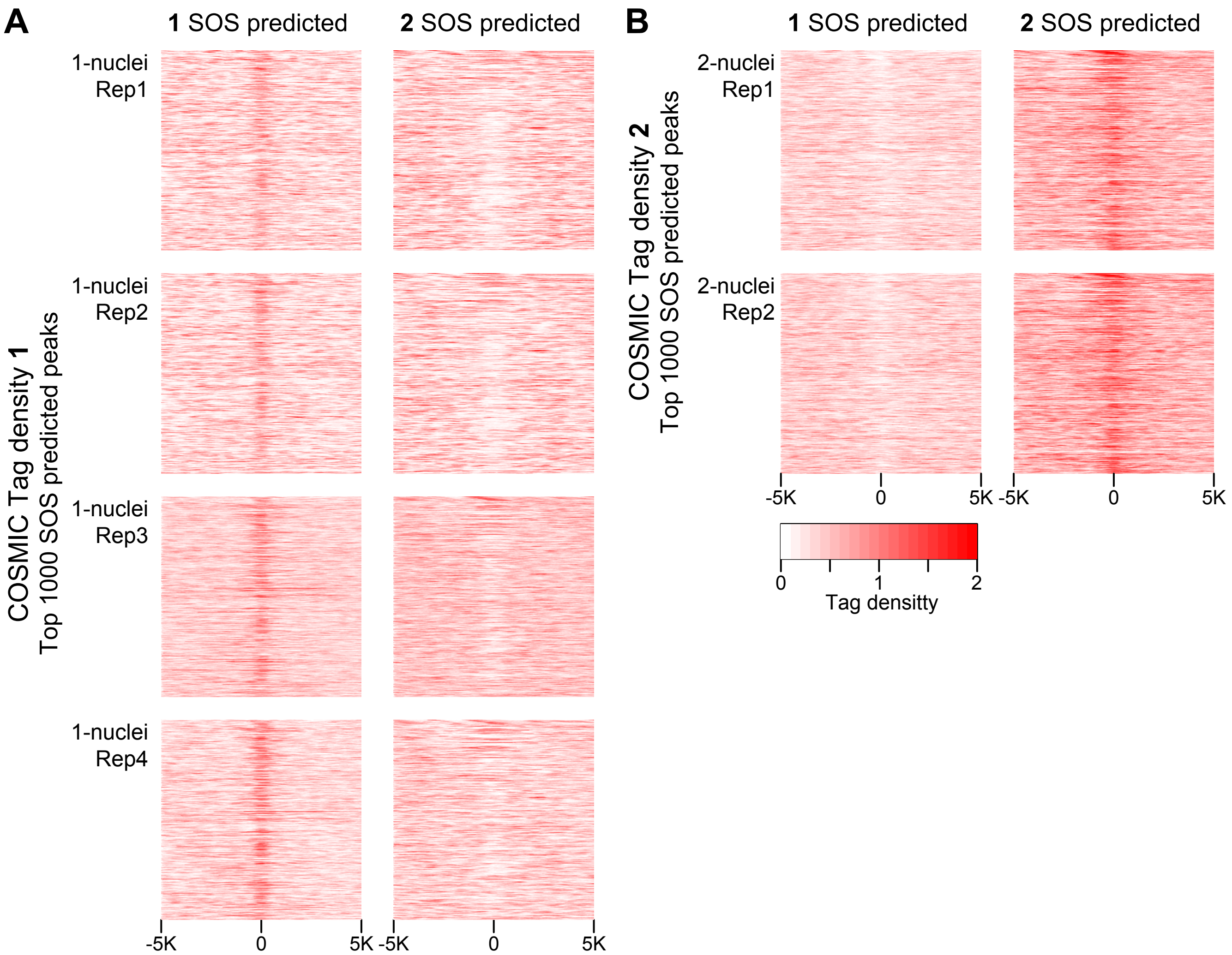
A
1 SOS predicted
2 SOS predicted
1-nuclei Rep1
1-nuclei Rep2
1-nuclei Rep3
1-nuclei Rep4
COSMIC Tag density 1
Top 1000 SOS predicted peaks
-5K
0
5K
-5K
0
5K
B
1 SOS predicted
2 SOS predicted
2-nuclei Rep1
2-nuclei Rep2
COSMIC Tag density 2
Top 1000 SOS predicted peaks
-5K
0
5K
-5K
0
5K
0
1
2
Tag densitty

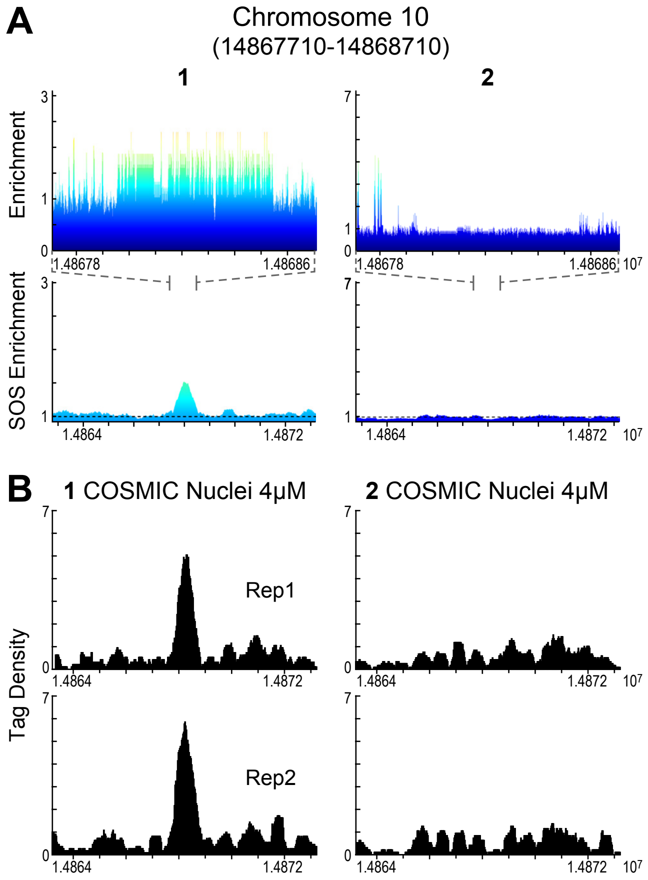
A
Chromosome 10
(14867710-14868710)
1
2
Enrichment
SOS Enrichment
1.48678
1.48686
$10^7$
1.4864
1.4872
B
1 COSMIC Nuclei 4μM
2 COSMIC Nuclei 4μM
Rep1
Rep2
Tag Density
