## Supplementary material for "Single position substitution of hairpin pyrrole-imidazole polyamides imparts distinct DNA-binding profiles across the human genome": S1 File

### SOS and GENOMESCAPE Heatmap at top 1000 COSMIC sites

### Contents


**1 1-nuclei-Rep1** **2**

**2 1-nuclei-Rep2** **3**

**3 1-nuclei-Rep3** **4**

**4 1-nuclei-Rep4** **5**

**5 2-nuclei-Rep1** **6**

**6 2-nuclei-Rep2** **7**

### 1 1-nuclei-Rep1

| CSI FILE | SOS heatmap | Genomescape |
|---|---|---|
| 1-RND1-50nM | 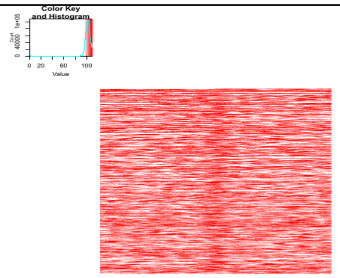  | 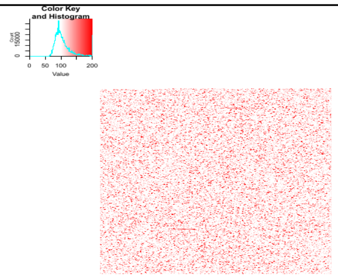  |
| 2-RND1-50nM | 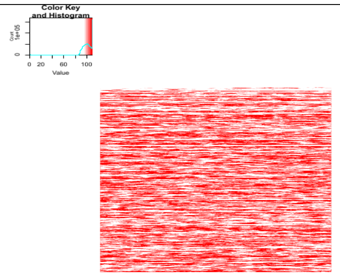  | 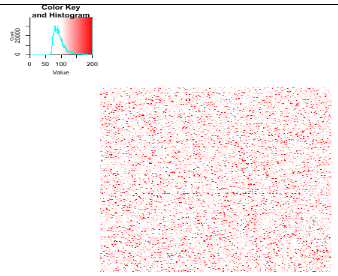  |

#### 2 1-nuclei-Rep2

| CSI FILE | SOS heatmap | Genomescape |
|---|---|---|
| 1-RND1-50nM | 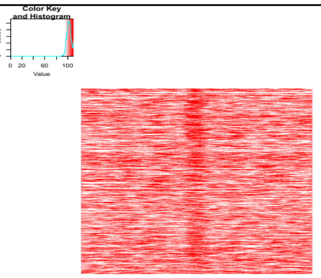  | 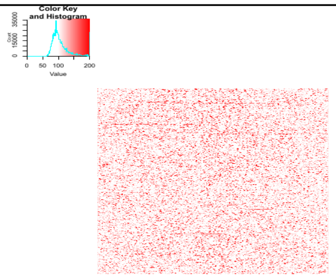  |
| 2-RND1-50nM | 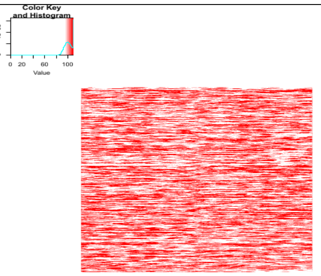  | 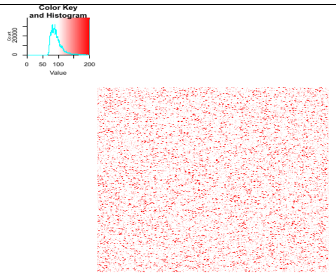  |

### 3 1-nuclei-Rep3

| CSI FILE | SOS heatmap | Genomescape |
|---|---|---|
| 1-RND1-50nM | 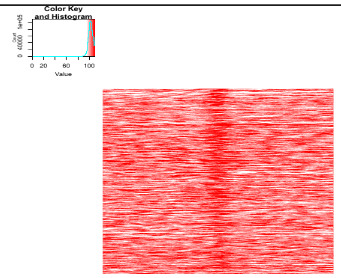 | 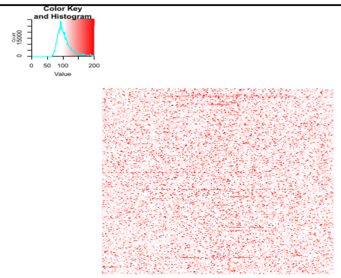 |
| 2-RND1-50nM | 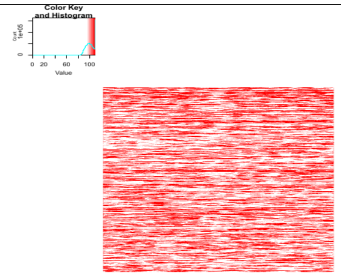 | 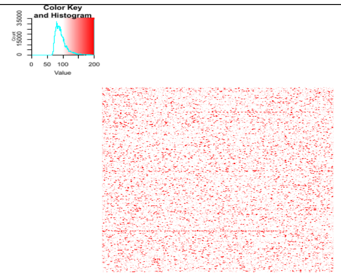 |

### 4 1-nuclei-Rep4

| CSI FILE | SOS heatmap | Genomescape |
|---|---|---|
| 1-RND1-50nM | 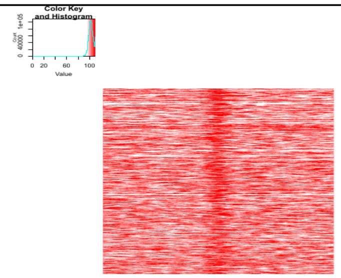  | 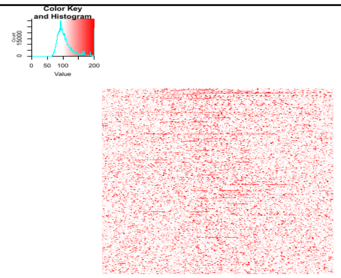  |
| 2-RND1-50nM | 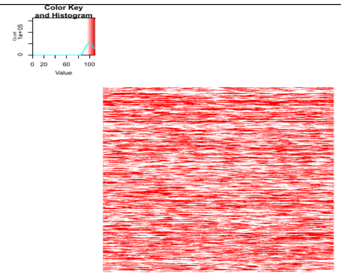  | 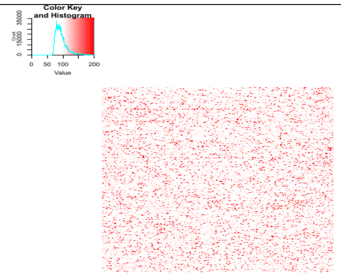  |

### 5 2-nuclei-Rep1

| CSI FILE | SOS heatmap | Genomescape |
|---|---|---|
| 1-RND1-50nM | 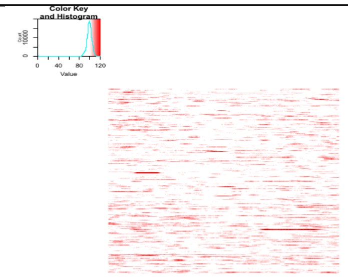 | 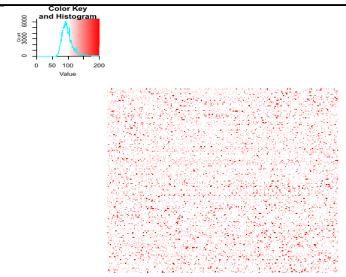 |
| 2-RND1-50nM | 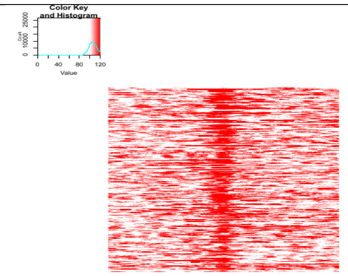 | 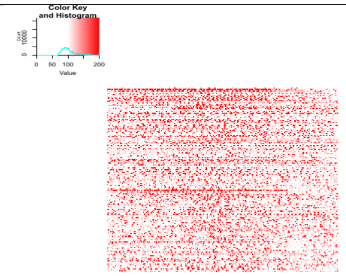 |

### 6 2-nuclei-Rep2

| CSI FILE | SOS heatmap | Genomescape |
|---|---|---|
| 1-RND1-50nM | 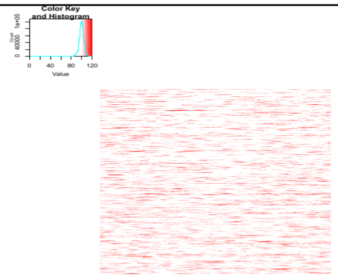 | 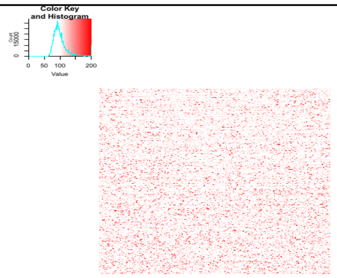 |
| 2-RND1-50nM | 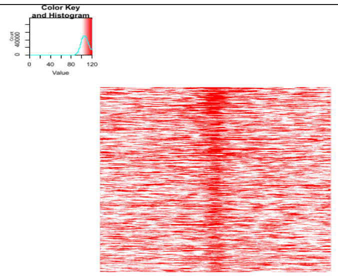 | 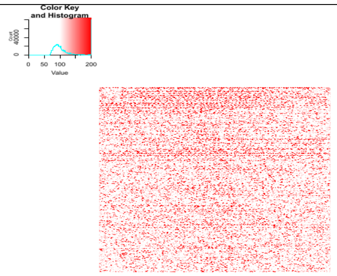 |
