## Supplementary material for "Single position substitution of hairpin pyrrole-imidazole polyamides imparts distinct DNA-binding profiles across the human genome": S1 File

### COSMIC tag density Heatmap at top 1000 SOS predicted sites

#### Contents


**1 WGWWCW-RND1-C50nM** **2**

**2 WGGWCW-RND1-C50nM** **3**

### 1 WGWWCW-RND1-C50nM

| COSMIC FILE | Tag density Heatmap |
|---|---|
| 1-nuclei-Rep1 | 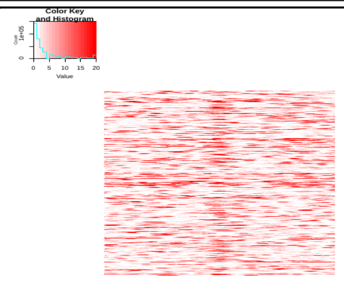 |
| 1-nuclei-Rep2 | 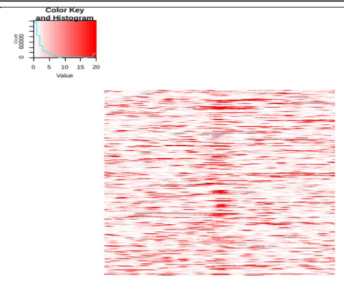 |
| 1-nuclei-Rep3 | 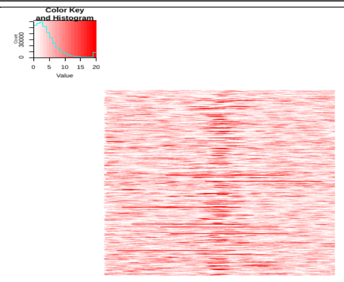 |
| 1-nuclei-Rep4 | 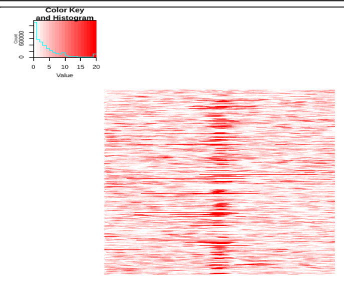 |
| 2-nuclei-Rep1 | 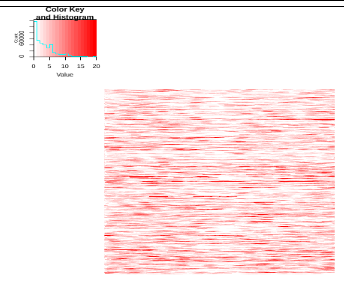 |
| 2-nuclei-Rep2 | 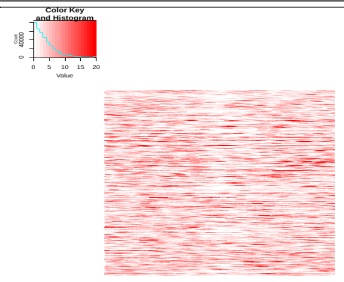 |

#### 2 WGGWCW-RND1-C50nM

| COSMIC FILE | Tag density Heatmap |
|---|---|
| 1-nuclei-Rep1 | 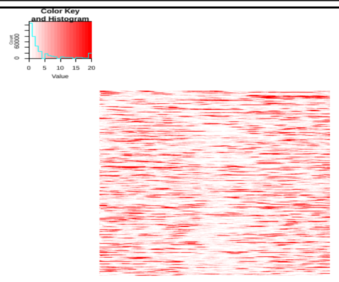 |
| 1-nuclei-Rep2 | 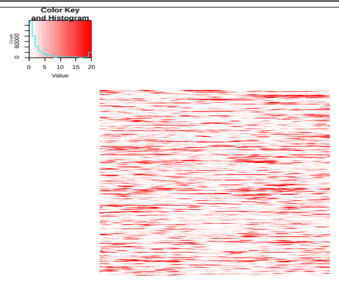 |
| 1-nuclei-Rep3 | 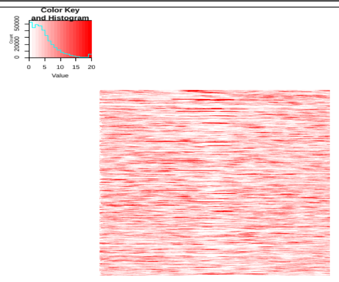 |
| 1-nuclei-Rep4 | 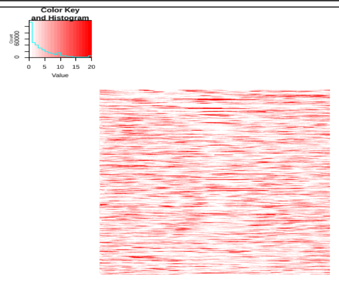 |
| 2-nuclei-Rep1 | 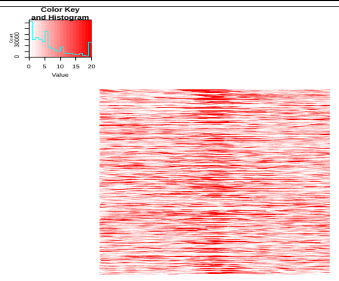 |
| 2-nuclei-Rep2 | 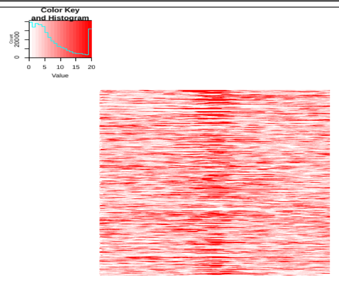 |
